# A Planar Switching Neuron Shares Logarithmic First-Spike Latency with Its Three-Dimensional Parent But Not Its Onset Mechanism

**DOI:** 10.64898/2026.10.17.759830

**Authors:** Giorgio Sonnino, Vadim Shlyonskiy, Thomas Erneux, David Gall

## Abstract

Reducing an electronic neuron from three variables to two can preserve a familiar voltage trace while changing the underlying dynamics that create it. We examine this distinction in a piecewise-linear planar reformulation of the three-dimensional neurosimilator. The planar model produces repetitive spike-like excursions without an imposed reset and admits an exact linear solution inside each of six switching regions. Direct comparison reveals only partial agreement between the two models. Although both generate large voltage excursions and an event rate that rises with applied drive, their waveforms and frequency-current gains differ substantially. A reanalysis of the three-dimensional, near-threshold data also corrects an earlier interpretation: over the resolved interval, the first-spike latency depends logarithmically on the normalized distance from threshold, matching the behavior of the planar model. However, this common functional form does not imply a shared onset mechanism. The three-dimensional equilibrium branch reaches a saddle-node limit point, whereas the planar equilibrium remains a stable node until it exits its admissible region at a switching boundary. While a stagewise analytical reconstruction captures individual voltage excursions, crossing-time errors accumulate unless Newton corrections are applied. The reduced system is therefore a useful electronic neuron model with transparent switching dynamics, rather than a dynamically equivalent copy of its three-dimensional parent.

## 1 Introduction

Low-dimensional neuron models are valuable because they isolate mechanisms that often disappear inside large, conductance-based systems. Beyond their role in elucidating neuronal dynamics, low-dimensional models can inform the development of neuromorphic circuits and their associated computational algorithms, with the prospect of more energy-efficient computational systems. Tight coupling of memory and computation, sparse event-based communication, and local learning offer routes to reducing data-transfer costs and supporting low-latency, real-time processing. These capabilities are particularly relevant to embedded applications constrained by size, weight, and power, while also offering potential benefits for large-scale computation [1]. Understanding which dynamical properties a reduced model preserves is therefore relevant to both theoretical analysis and hardware design. However, this economy carries a risk. Two models can produce similar voltage traces and still organize threshold crossing, recovery, and repetitive activity in fundamentally different ways; a reduction should therefore be judged by more than the mere appearance of a spike. This issue is especially apparent in analog electronic neurons. Their state variables arise from physical circuit elements, and their thresholds arise from switching components, meaning the correspondence with ion-channel kinetics is functional rather than microscopic. Classical reductions like the FitzHugh-Nagumo and McKean systems were built to retain specific dynamical features of excitability [2, 3]. Piecewise-linear models extend that approach, allowing exact calculations between switching events [4, 5]. More recent circuit designs leverage threshold devices, memristive elements, and coupled membranes to achieve a broad range of firing patterns [6-8]. Their physical implementations make a basic question unavoidable: which properties belong to the observed waveform, and which belong to the underlying phase-space geometry? The three-dimensional Neurosimilator studied in Ref. [9] combines smooth regenerative feedback with voltage-controlled switching, supporting periodic, quasiperiodic, and chaotic responses alongside a direct analog realization. The two-variable framework analyzed here is the planar model developed in the companion letter [10], where an instantaneous, threshold-activated term replaces the fast depolarizing state. This is a heuristic reformulation rather than an asymptotic reduction of the three-dimensional equations or a reduction of Hodgkin-Huxley channel kinetics [11, 12]. The present Letter asks one focused question: what survives this reformulation? We compare the waveform, the frequency-current relation, and the geometry of firing onset. In [9], starting from the analytically derived current limit point*I*_*LP*_ = 0.296 *V ms*^*−*1^, we applied the inverse square root law to the repetitive spiking period.

In this letter, we also analyze the gap spanning the interval from the onset of the critical (rheobasic) current *I*_*c*_ = 0, 208 *V ms*^*−*1^ to *I*_*LP*_ = 0.296 *V ms*^*−*1^ for both 3D and 2D models. Furthermore, we test the analytical timing construction against event-resolved integration. The comparison resolves an apparent contradiction: the first-spike latency of both models, 3D and 2D, follows a logarithmic law over the measured interval^1^, yet the mechanisms remain different. Shared scaling, by itself, does not establish dynamical equivalence.

## 2 Planar switching model

The reduced model tracks a voltage-like variable *v* and a recovery variable *k*:

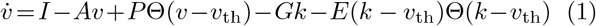

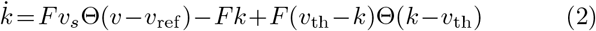

Here, Θ denotes the Heaviside step function and *P* = *F* (*v*_*s*_ − *V*_0_), where *V*_0_ = 1 V serves as the circuit reference. The input *I* represents the injected current normalized by membrane capacitance. Because of this scaling, its unit is voltage per unit time rather than current. Throughout this study, we fix the parameters as:

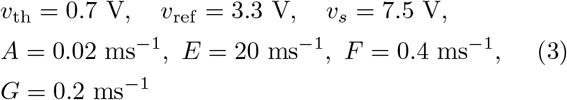

These mathematical terms map directly to circuit-level leakage, threshold-activated depolarization, recovery, and repolarization; they are not intended to represent specific physiological sodium or potassium currents. Crucially, Equations (**1**) and (**2**) do not rely on an imposed reset mechanism. Instead, the continuous vector field naturally routes the trajectory through a large depolarizing excursion, followed by repolarization and subsequent recovery. The boundaries defined by *v* = *v*_th_, *v* = *v*_ref_, and *k* = *v*_th_ slice the phase plane into six distinct linear regions, where a standard firing cycle steps sequentially through:

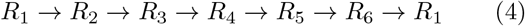

## 3 What the reduction preserves

A direct comparison between the parent 3D model [9] and the planar model introduced in the companion letter [10] is illustrated in Figure 1. Because the individual voltage traces in panel (A) are normalized separately, this panel evaluates waveform morphology and timing rather than physical amplitude. While both traces display a pronounced excursion followed by a recovery phase, the similarities end there; the 2D event is visibly sharper, and their recovery trajectories operate on distinct timescales.

**Figure 1:**
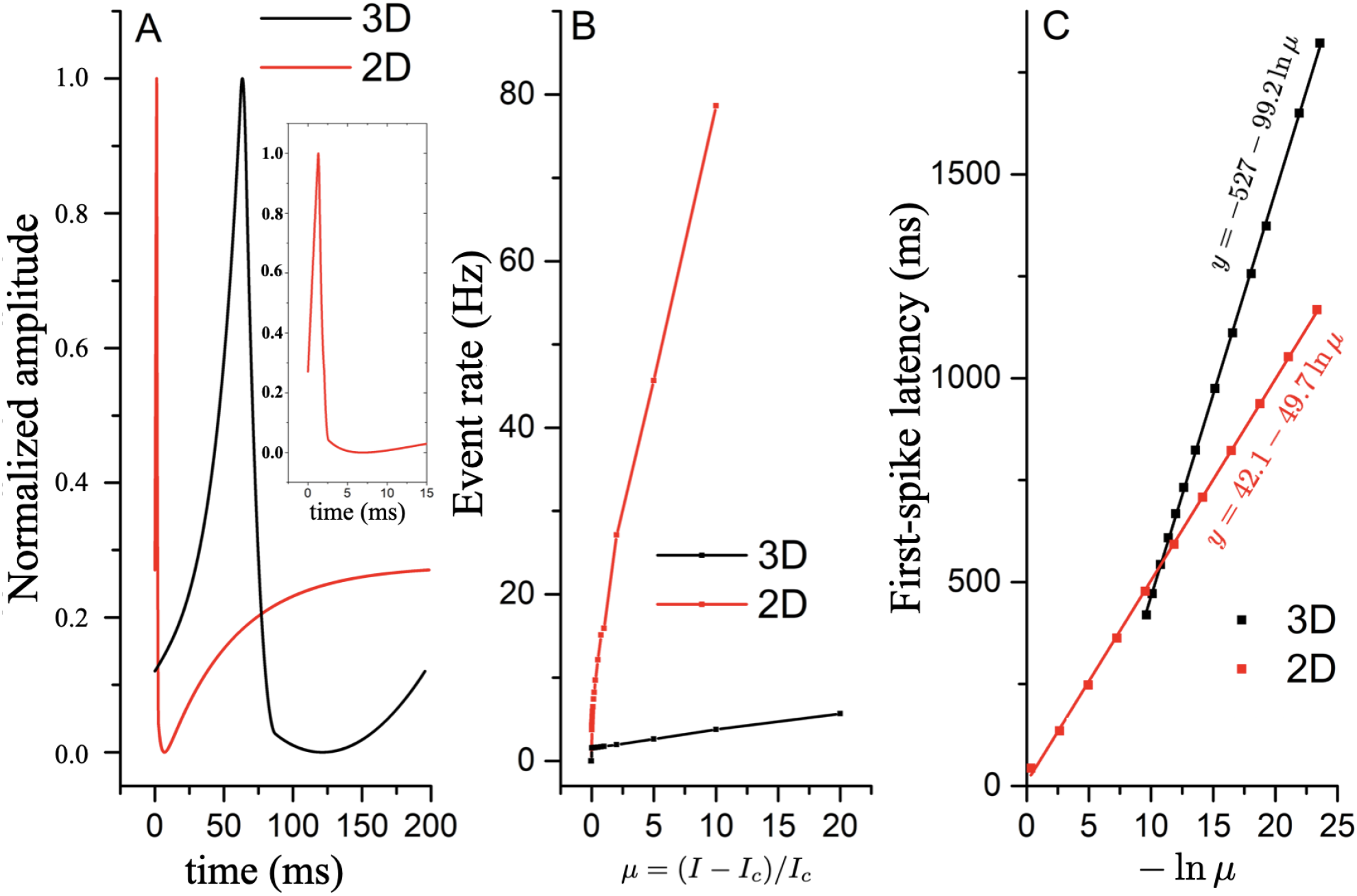
Comparison of the 3D and 2D models. (A) Normalized voltage amplitude versus time. The inset resolves the sharp 2D excursion over 0–15 ms. The 3D trace reaches its maximum later and follows a different recovery path. (B) Event rate as a function of the normalized drive *µ* = (*I − I*_*c*_)*/I*_*c*_. Both curves rise with input, but the 2D response has a much larger gain over the plotted range. (C) Spiking period versus *−* ln *µ*. The straight lines are 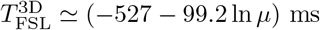 and 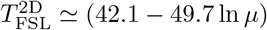 over the intervals shown. Black and red denote the 3D and 2D models, respectively.

To establish a dimensionless metric for the applied drive, we define the relative distance-from-threshold as:

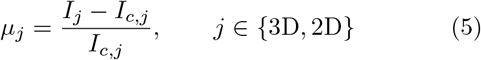

where *I*_*c*,3D_ = *I*_LP_ represents the three-dimensional limit-point input and *I*_*c*,2D_ = *I*_th_ = *Av*_th_ denotes the planar switching threshold. Note that these two reference values carry distinct physical meanings; Equation (**5**) simply maps both inputs onto a comparative scale based on their proximity to threshold. Panel (B) plots the resulting event rate under constant input. Although both rates scale upward with *µ*, their gains diverge by more than an order of magnitude across the illustrated ranges. Specifically, the 2D model approaches 80 Hz while the 3D model stays below roughly 7 Hz. Ultimately, the reduction captures the directional trend of the input-output response, rather than its exact quantitative gain. Panel (C) shifts focus to the spiking period rather than the mean interspike interval. Assuming a constant input step initiates at *t* = *t*_step_ and the initial upward threshold crossing happens at *t* = *t*_1_, the latency is given by:

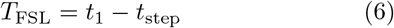

Isolating the first spike is critical in the 3D system. Following the initial spike, the state-space trajectory re-enters the bottleneck from an entirely different region, making subsequent interspike intervals substantially shorter. Averaging these subsequent cycles with the initial latency would conflate distinct types of state-space transits. Such spiking behavior is not unique and is also known for the theta-neuron model and for some parameter sets in the Hodgkin-Huxley model [13]. We note that Ref. [9] previously modeled the repetitive spiking period starting from the analytically derived current limit point *I*_*LP*_ = 0.296 *V ms*^*−*1^ in the 3D data using an inverse-square-root scaling law. However, a reanalysis of this regime challenges that classification. Across the resolved interval, the 3D data vary linearly with −ln *µ*, mirroring the behavior of the 2D data. The fitted slopes do differ, yielding approximately 99.2 ms for the 3D model and 49.7 ms for the 2D model. We treat this relationship strictly as an empirical scaling valid over the observed range. Consequently, we do not extrapolate these fits to deduce the asymptotic normal form of the 3D limit point, nor do we attempt to infer a zero steady-state firing frequency solely from a first-spike metric.

## 4 The same latency law can arise from different geometry

Beneath the firing threshold, every Heaviside term drops out, leaving the planar equilibrium at

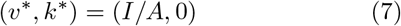

The corresponding Jacobian eigenvalues are simply *A* and −*F*. Because both values remain strictly negative as *I* scales upward, the system experiences no loss of stability in the traditional sense. Rather, at the critical input level

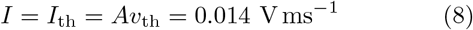

the stable node encounters *v* = *v*_th_, which defines the structural boundary where Eq. (7) ceases to be valid. Once *I > I*_th_, the virtual equilibrium *v*^*\**^ = *I/A* moves outside this domain, forcing the trajectory into the adjacent linear subsystem. Crucially, no pair of planar fixed points collides, and neither eigenvalue passes through zero. The transition is fundamentally a boundary-crossing event rather than a saddle-node bifurcation of the underlying planar vector field. This specific geometry is illustrated clearly in Figure 2, which also provides the physical intuition behind the planar model’s logarithmic latency behavior. Throughout the prolonged subthreshold interval spent within *R*_1_, the variable *k* relaxes to a value very close to zero, simplifying the voltage dynamics to

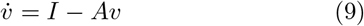

**Figure 2:**
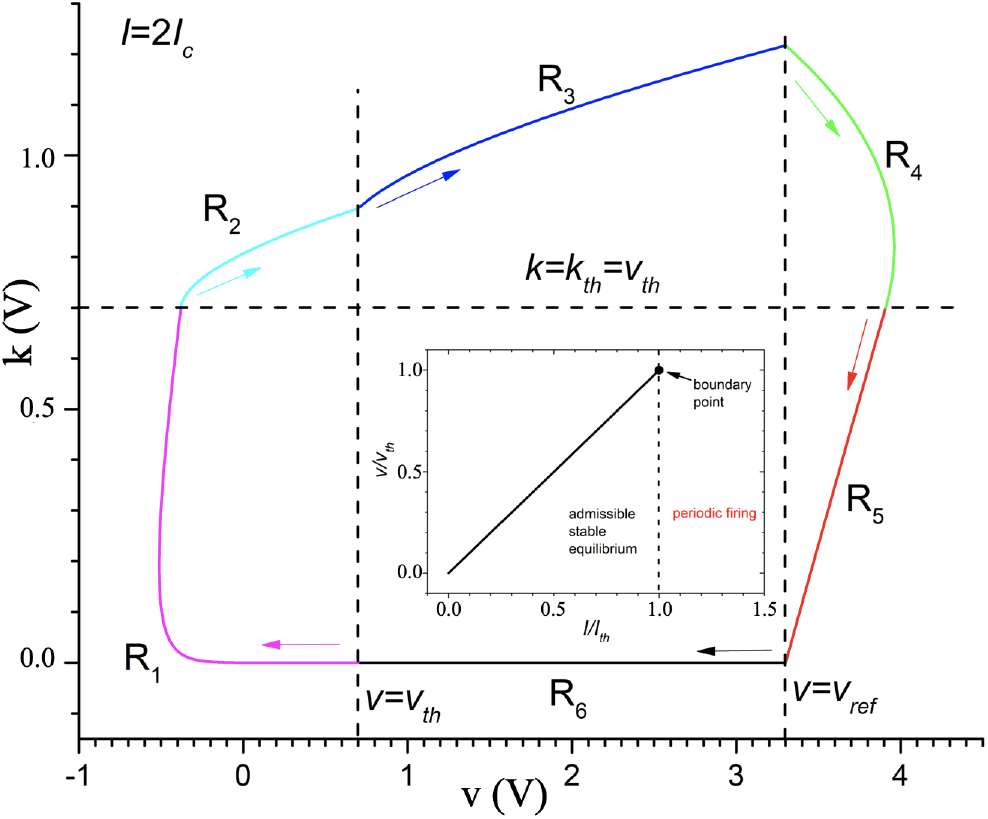
Trajectory through the six switching regions of the 2D model. The phase-plane orbit at *I* = 2*I*_*c*_ follows the sequence in Eq. (4); arrows indicate the direction of motion. Dashed lines mark *v* = *v*_th_, *v* = *v*_ref_, and *k* = *v*_th_. The inset shows the admissible stable equilibrium below *I*_th_ and its arrival at the switching boundary. Fixed-step RK4 provides uniformly spaced plotting data but needs a small step near a switch. Event-resolved DOP853 locates boundary crossings accurately; its nonuniform internal nodes were not used as a plotting grid.

Integrating this relation from an initial condition *v*_0_ reveals that the time needed to strike the *v*_th_ threshold is

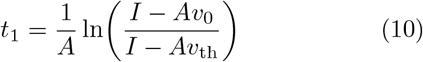

rovided *v*_0_ stays well insulated from *v*_th_ in the limit 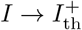, the leading-order behavior scales as

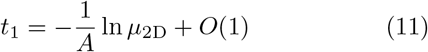

This derivation predicts a scaling coefficient of 1*/A* = 50 ms, striking an excellent match with the empirically fitted value of 49.7 ms shown back in Fig. 1(C). The logarithmic trend is thus a direct consequence of a slow drift toward a switching manifold. Remarkably, the 3D model yields an identical empirical scaling over the tested regime, even though its equilibrium branch vanishes via a true saddle-node limit point. This warns that a shared functional scaling in a single macroscopic observable can easily obscure deeply contrasting internal bifurcation structures.

## 5 Accuracy of the stagewise timing calculation

Within any given region *R*_*j*_, the local dynamics obey a linear system of the form

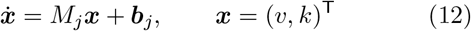

yielding an analytical trajectory expressed as:

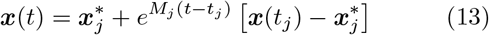

Determining the exit time is where the algebraic challenge arises. Evaluating Eq. (13) at the subsequent boundary condition typically yields a transcendental sum of exponentials governed by differing decay rates. Consequently, while the intra-stage trajectory is exact, a closed-form expression for the exact manifold crossing time is generally out of reach. For this reason, we treat the reconstructed orbit as a stagewise analytical approximation rather than a true closed-form solution of the full periodic cycle (detailed expressions of the analytic equations can be found in [10]). Defining the crossing equation in region *R*_*j*_ as *H*_*j*_(*τ*_*j*_) = 0, an initial guess 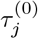 can be iteratively refined via Newton’s method:

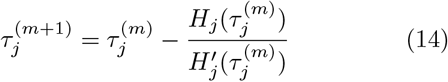

Because the final state of one linear segment serves as the initial condition for the next, even minor errors in individual *τ*_*j*_ values inevitably cascade into a macroscopic phase drift over successive cycles. Figure 3 directly quantifies this error accumulation. At driving strengths of 2*I*_th_ and 5*I*_th_, the uncorrected reconstruction successfully captures the primary excursion profile and repetitive rhythm; however, the simulated spike times exhibit a linear drift relative to the numerical baseline. The positive slopes indicate that the method systematically overpredicts the true period. Implementing a single Newton correction curtails this drift by roughly a factor of three, while a second iteration drops the error slope by approximately two orders of magnitude compared to the uncorrected baseline. This approach proves highly valuable for dissecting individual phase-space stages and tracking short-term timing properties, provided crossing corrections are strictly enforced. Conversely, uncorrected long-term trajectories are unreliable for diagnosing irregular spiking behavior or underlying bifurcations. As a numerical baseline, we integrated Eqs. (1) and (2) using an adaptive DOP853 routine coupled with explicit root-finding functions to pin down the three switching boundaries. This integration maintained a relative tolerance of 10^*−*11^, an absolute tolerance of 10^*−*13^, and a maximum allowable step size of 0.25 ms. To safeguard the display grid from artifacts of the solver’s internal adaptive stepping, dense output was generated and sampled on a separate, uniformly spaced plotting mesh.

**Figure 3:**
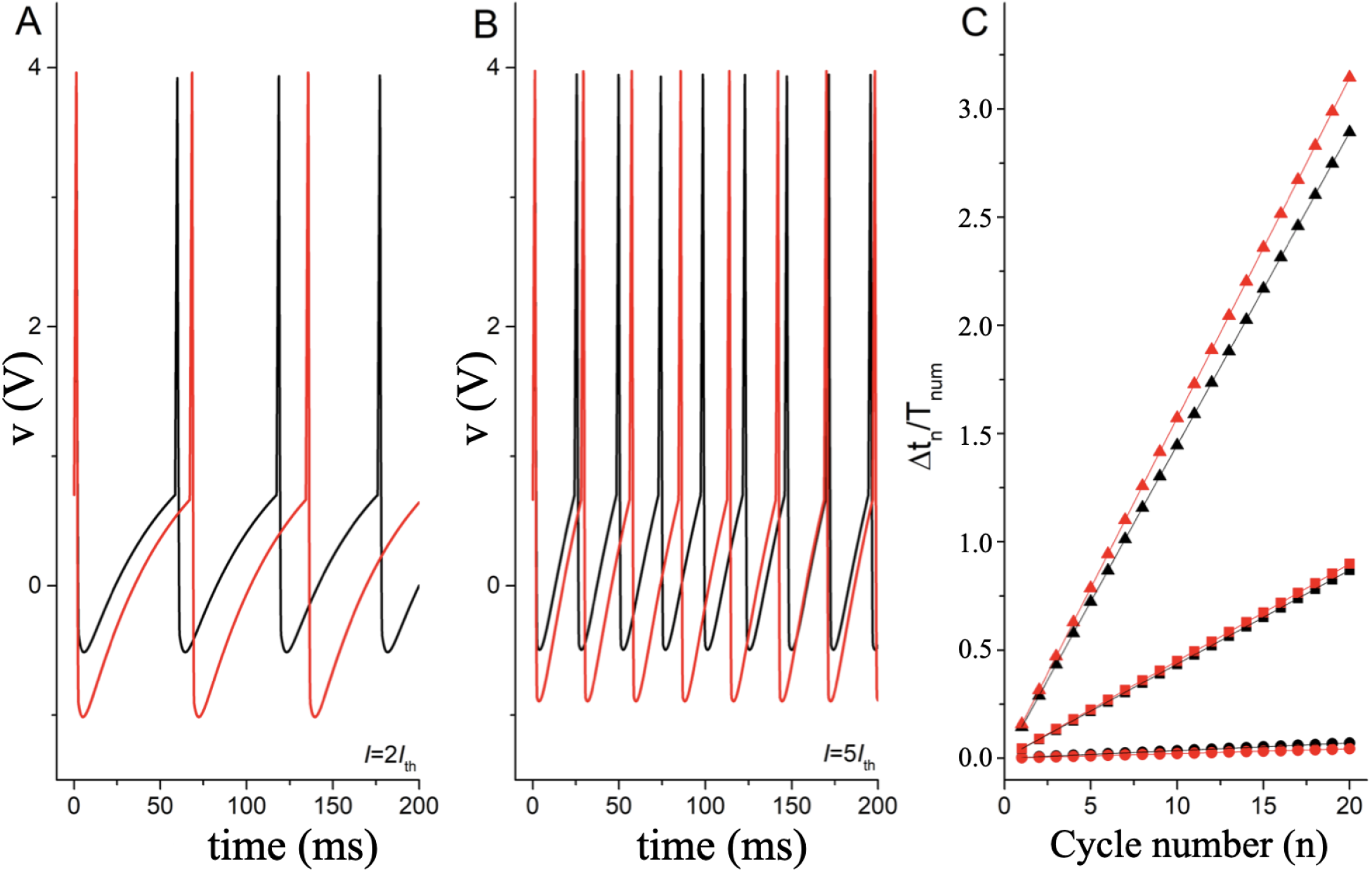
Accuracy of the six-stage analytical reconstruction. The stagewise approximation is compared with event-resolved DOP853 integration at (A) *I* = 2*I*_th_ and (B) *I* = 5*I*_th_. Black curves are numerical; red curves are the uncorrected reconstruction. (C) Normalized cumulative timing error Δ*t*_*n*_*/T*_num_, where 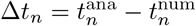. Black and red symbols correspond to 2*I*_th_ and 5*I*_th_. Triangles show the uncorrected approximation, with slopes 0.1445 and 0.1571; squares show one Newton correction, with slopes 0.0435 and 0.0449; circles show two corrections, with slopes 0.00352 and 0.00217, for black and red data, respectively. The numerical reference used rtol = 10^*−*11^, atol = 10^*−*13^, and Δ*t*_max_ = 0.25 ms.

## 6 Circuit realization, limits, and interpretation

The planar dynamical equations map directly onto the compact hardware platform illustrated in Fig. 4. Threshold tracking and recovery mechanisms are handled by discrete switching and comparator stages, with the entire printed circuit board powered via a standard USB-C interface. While this physical implementation demonstrates that the two-variable switching framework is readily realizable using off-the-shelf electronic components, it serves as a functional analog; it does not map the underlying mathematical terms onto literal biophysical ion currents. The circuit schematic of the 2D model can be found in the companion manuscript [10] and in [14]. The signal provided by the electronic circuit has been compared with the numerical and stepwise closed-form solutions of the ordinary differential equations (ODEs) (see Figure 5). Several scope boundaries constrain the claims made in this work. Foremost, the dimensional reduction is fundamentally heuristic: it is engineered to capture specific input-output response characteristics, but it sacrifices the true 3D bifurcation topology, the quantitative frequency-current gain profile, and the broader spectrum of complex periodic, quasiperiodic, or chaotic behaviors. Furthermore, because the model relies heavily on multiple Heaviside step functions, precise numerical event localization becomes paramount. Utilizing a coarse, fixed integration step introduces localized boundary-crossing shifts that can rapidly degrade the integrity of simulated spike timings. An additional limitation stems from the fact that the piecewise analytical calculations remain exact only within the interior of each linear domain. Approximations made when solving for crossing times inevitably inject phase errors that accumulate over time, an issue that becomes particularly pronounced under high-drive conditions. Similarly, the logarithmic first-spike latency observed in the 3D data must be treated strictly as an empirical fit over the investigated regime, rather than a rigorous diagnostic of the asymptotic normal form governing the saddle-node bifurcation. Finally, shedding the third state fundamentally strips the system of its intrinsic memory. Consider the linearized dynamics of this missing variable:

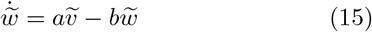

Following the decay of initial transients, this state evolves according to:

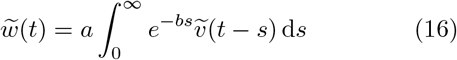

**Figure 4:**
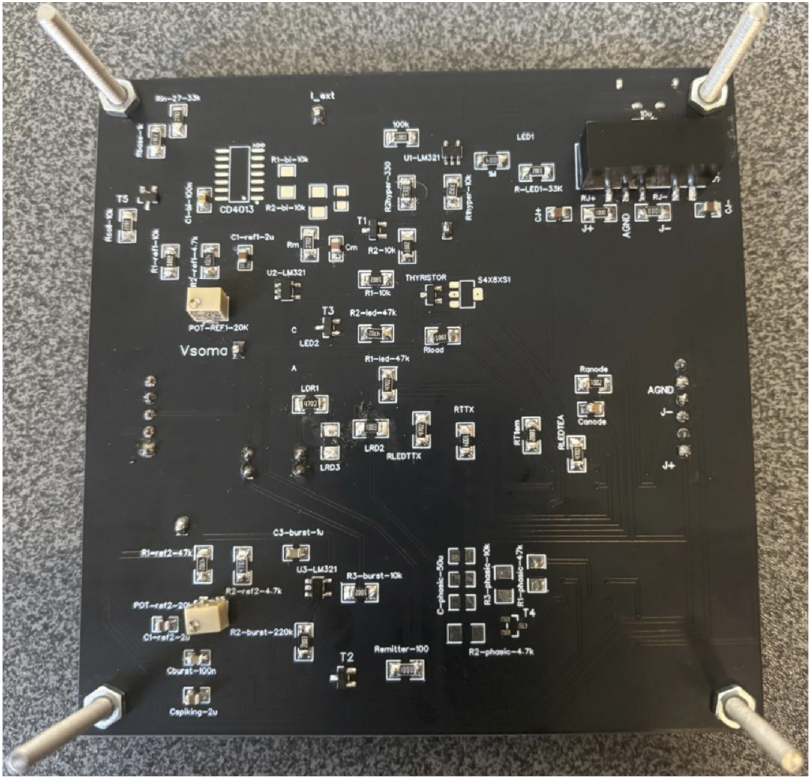
Printed-circuit realization of the planar electronic neuron. The board contains the switching and recovery stages associated with Eqs. (1) and (2). Power is supplied through USB-C, with the required rails generated on board.

**Figure 5:**
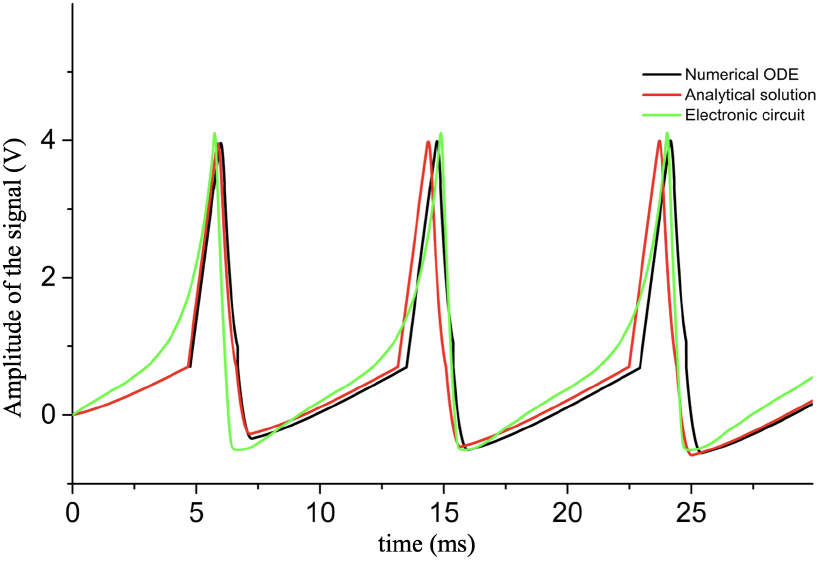
Piecewise-linear neuron model with analytical and hardware realisation. This figure compares the numerical solution of the ODEs (in blue), the stepwise closed-form solution (in red), and the signal provided by the electronic circuit shown in Figure 4.

An instantaneous, state-dependent threshold function is mathematically incapable of replicating this type of distributed temporal memory. Although incorporating delayed or kernel-based feedback mechanisms offers a compelling direction for future inquiry, doing so would yield a two-variable model embedded with a historical continuum, rather than a strictly two-dimensional autonomous dynamical system. We intentionally leave such extensions outside the scope of the present study.

## 7 Conclusion

The planar model reproduces three prominent features of its 3D parent: a large spike-like excursion, a drive-dependent event rate, and a near-threshold first-spike latency that scales logarithmically over the measured interval. However, a quantitative comparison exposes exactly what is lost in the reduction. Waveform timing alters, the frequency–current gain shifts sharply, and the richer behaviors of the full 3D dynamics are absent. The central result is geometrical. In the planar model, a stable node reaches the boundary of its admissible region, forcing the trajectory into a six-stage switching cycle without encountering a planar saddle-node bifurcation. By contrast, the 3D model reaches a genuine saddle-node limit point. Consequently, their first-spike latency data can share an identical empirical logarithmic form without sharing the same onset mechanism. The reduction does offer a practical advantage. Each segment of the trajectory remains analytically transparent, and applying just two Newton corrections keeps the accumulated timing error small over a tested twenty-cycle window. The resulting system is therefore best described as a standalone two-variable piecewise-linear electronic neuron with a well-defined operating range, rather than a dynamically equivalent reduction of the 3D Neurosimilator.

## CRediT authorship contribution statement

The authors contributed equally to the work.

## Data and code availability

The code used to generate and analyse the results is available at https://gitlab.ulb.be/vshlyons/2d-neurosimilator. PCB photos of the 3D model circuit can be seen in [9] and [14].

## Footnotes

1 Notably, Ref. [9] concluded that starting from the analytically derived current limit point *I*_*LP*_ = 0.296 *V ms*^*−*1^, the repetitive spiking period (RSP) obeys an inverse-square-root law. The inverse-square-root dependence provides a reasonably good description of the data, with *R*^2^ ≃ 0.95, and therefore cannot be regarded as incompatible with the observations based on goodness of fit alone. However, the logarithmic dependence yields a substantially closer fit *R*^2^ ≥ 0.99. We therefore favor the logarithmic representation empirically.

